# CaMKIIα promoter-controlled circuit manipulations target both pyramidal cells and inhibitory interneurons in cortical networks

**DOI:** 10.1101/2022.06.08.495358

**Authors:** Judit M. Veres, Tibor Andrasi, Petra Nagy-Pal, Norbert Hajos

**Affiliations:** ELRN Institute of Experimental Medicine, Budapest, Hungary; The Linda and Jack Gill Center for Molecular Bioscience, Indiana University Bloomington, Indiana, USA; János Szentágothai School of Neurosciences, Semmelweis University, Budapest, Hungary; Program in Neuroscience, Department of Psychological and Brain Sciences, Indiana University Bloomington, Indiana, USA

**Author notes:** These authors equally contributed to the study (Judit M. Veres, Tibor Andrasi). **Corresponding author:** Norbert Hajos, Indiana University Bloomington.

## Abstract

A key assumption in studies of cortical functions is that excitatory principal neurons, but not inhibitory cells express calcium/calmodulin-dependent protein kinase II subunit α (CaMKIIα) resulting in a widespread use of CaMKIIα promoter-driven protein expression for principal cell manipulation and monitoring their activities. Using neuroanatomical and electrophysiological methods we demonstrate that in addition to pyramidal neurons, multiple types of cortical GABAegic cells are targeted by adeno-associated viral vector (AAV) carrying the CaMKIIα-Channelrhodopsin 2-mCherry construct. We show that the reporter protein, mCherry can visualize a large fraction of different interneuron types, including parvalbumin (PV), somatostatin (SST), neuronal nitric oxide synthase (nNOS) and neuropeptide Y (NPY)-containing GABAergic cells, which altogether cover around 50% of the whole inhibitory cell population in cortical structures. Importantly, the expression of the excitatory opsin Channelrhodopsin 2 in the interneurons effectively drive spiking of infected GABAergic cells even if the detectability of reporter proteins is ambiguous. Thus, our results challenge the use of CaMKIIα promoter-driven protein expression as a selective tool in targeting cortical glutamatergic neurons using viral vectors.

## Introduction

Cortical structures comprise excitatory principal cells and various types of inhibitory neurons, including PV-containing interneurons responsible for perisomatic inhibition, SST-expressing dendritic inhibitory cells, nNOS-containing GABAergic cells, a majority of which project to other brain regions and NPY-expressing interneurons giving rise to slow synaptic inhibition in the dendritic compartments (Tremblay *et al*., 2016; Pelkey *et al*., 2017; Hajos, 2021). To understand the neuronal network operation at cellular and circuit levels, controlling or monitoring the activity of defined neuronal populations selectively have become a gold standard method in neuroscience research (Bernstein *et al*., 2012; Sternson & Roth, 2014; Kim *et al*., 2017; Luo *et al*., 2018). This goal can be achieved by using genetically modified mice, viral vectors carrying constructs that induce protein expression specifically in a cell group, or combination of these two methods (Fenno *et al*., 2014; Huang, 2014; Song & Palmiter, 2018). To manipulate the glutamatergic principal neurons in cortical networks, CaMKIIα promoter-driven expression of various effector proteins is a usual choice (Basu *et al*., 2008; Scheyltjens *et al*., 2015; Egashira *et al*., 2018). This widespread approach relies on the results obtained by immunohistochemical studies showing that CaMKIIα is present in cortical glutamatergic cells, but not in GABAergic interneurons (Liu & Jones, 1996; Sik *et al*., 1998).

In contrast to these examinations using immunostaining, some studies reported that the CaMKIIα promoter either using CaMKIIα-Cre mice or viral vectors carrying CaMKIIα-GFP also visualized GABAergic interneurons in cortical areas in addition to glutamatergic pyramidal neurons (Watakabe *et al*., 2015; Radhiyanti *et al*., 2021). Moreover, a recent study using microRNA-guided neuron tagging has labelled PV and SST interneurons in the motor cortex in a CaMKIIα promoter-dependent manner (Keaveney *et al*., 2020). These results have already raised the concern whether the use of CaMKIIα promoter is eligible for specific targeting of glutamatergic excitatory neurons in cortical structures. However, no systematic investigation has been carried out to clarify the prevalence of CaMKIIα promoter-controlled expression of proteins in different GABAergic cell types and whether the expression has any functional relevance in inhibitory neurons that would limit the use of the CaMKIIα promoter in cortical areas, including neocortex, hippocampus and basolateral amygdala complex.

To address these critical questions for circuit studies, we first assessed the ratio of neurons that express the reporter protein mCherry under the control of CaMKIIα promoter in four distinct interneuron types, in three different cortical regions. Next, we directly tested the functional consequence of CaMKIIα promoter-driven expression of Channelrhodopsin 2 (ChR2) in interneurons. We found that the reporter protein expression in GABAergic cells under CaMKIIα promoter showed a cell type-and region-dependent variability. In line with these results, light activation of ChR2 in GABAergic cells reliably depolarized and drove their spiking, even in those interneurons in which the level of the reporter proteins tested by immunolabeling was below the detection threshold.

## Results

We first investigated the ratio of GABAergic cells in which CaMKIIα promoter drives protein expressions using viral vectors. To this end, we injected AAV5-CaMKIIα-ChR2-mCherry into the CA1 region of the hippocampus, medial prefrontal cortex (mPFC) and basolateral amygdala (BLA) in wild type mice. After 4-5 weeks of injection, we performed immunostaining on fixed tissues containing the virus infected areas. The presence of mCherry was investigated in inhibitory cells immunolabeled for neurochemical markers typically expressed by distinct, largely non-overlapping interneuron groups. We observed that in the mPFC and BLA, mCherry signal could be detected in as high as 80% of interneurons immunostained for PV. In contrast, there was no colocalization between the mCherry and PV immunolabeling in the hippocampus (Fig. 1A, B). In SST-expressing interneurons, mCherry content varied in the examined structures. The largest fraction of colocalization was found in the BLA (79%), whereas the lowest in the hippocampus (9%, Fig. 1C, D). For nNOS-expressing GABAergic cells we found that the highest ratio of mCherry co-labeled cells was in the BLA (66%) in comparison to those observed in the hippocampus and mPFC (Fig. 1E, F). Lastly, mCherry content was obvious in at least half of interneurons expressing NPY in all cortical areas investigated (Fig. 1G, H). These results clearly show that i) at least four distinct group of GABAergic inhibitory cells can be infected by the use of CaMKIIα promoter and ii) the fraction of inhibitory cells expressing mCherry under the control of this promoter varies between cortical structures, but it can be as high as 80% in a given interneuron population. Thus, in addition to glutamatergic principal neurons, a significant number of GABAergic cells are targeted by CaMKIIα promoter in the three cortical areas examined.

**Fig. 1.**
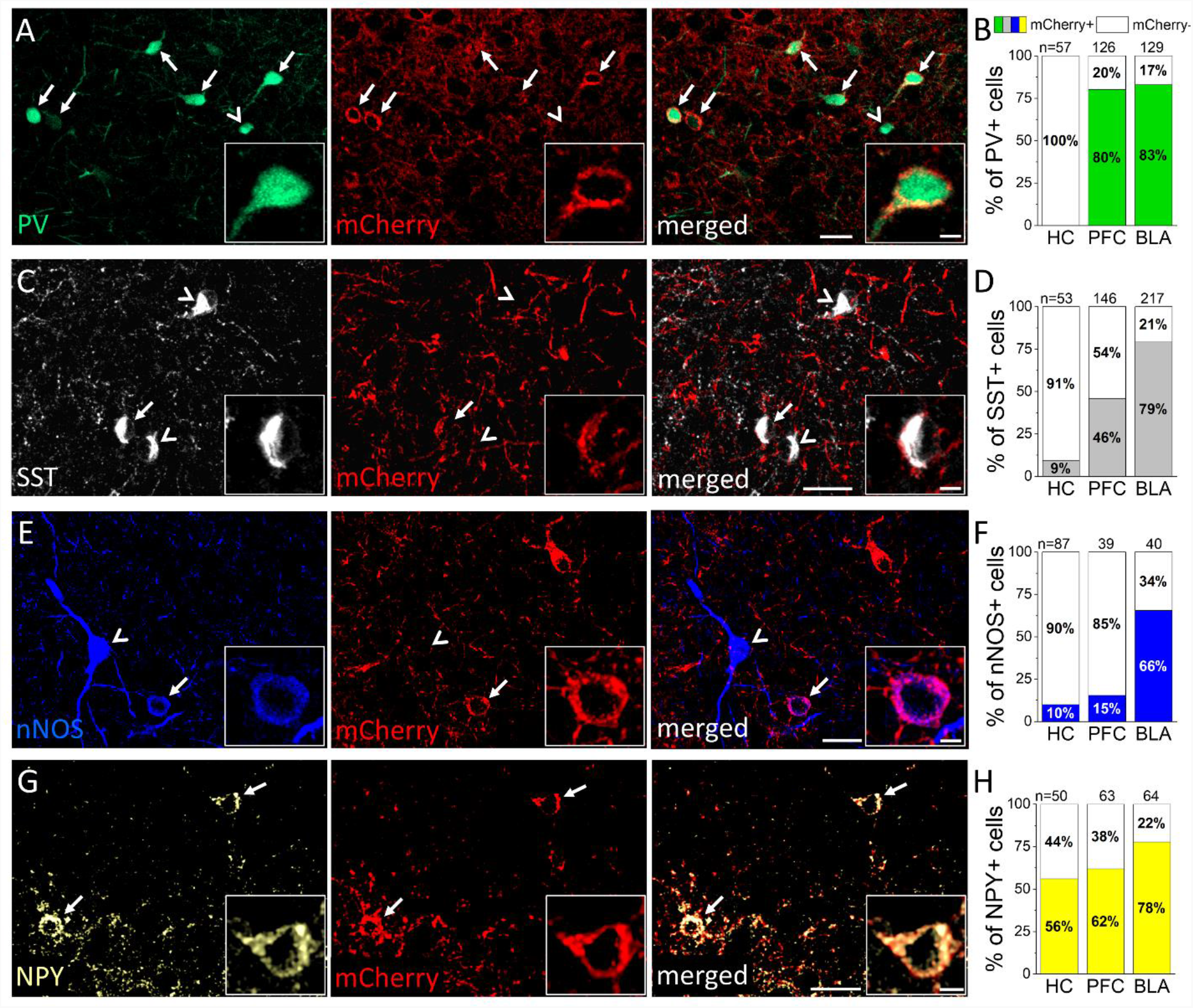
CaMKIIα promoter-driven expression of the red fluorescent protein mCherry visualizes a substantial portion of various interneuron types in three cortical areas. **(A, C, E, G)** Confocal microscopic images taken from the basolateral amygdala (A, C, E) and medial prefrontal cortex (G) of fluorescently immunolabeled parvalbumin (PV, *green*), somatostatin (SST, *white*), neuronal nitric oxide synthase (nNOS, *blue*) and neuropeptide Y (NPY, *yellow*) containing interneurons, respectively, together with mCherry (*red*) expression in neurons infecting by AAV5-CaMKIIα-ChR2-mCherry construct. Arrows label mCherry containing interneurons, arrowheads label those lacking mCherry signal. Scale bars, 25 µm and 5 µm (insets). (**B, D, F, H**) Ratio of mCherry expression in the different interneuron types in the hippocampus (HC), medial prefrontal cortex (PFC) and basolateral amygdala (BLA) regions. n=3 mice, 36 slices, number of examined cells in a given region is indicated above each bar.

In the next set of experiments, we tested whether the CaMKIIα promoter-driven expression of ChR2 linked to the mCherry can affect the firing of distinct types of inhibitory cells. We prepared acute slices from brain regions where viral vectors were injected and performed targeted recordings from GFP-or EYFP-expressing GABAergic cells. To isolate the direct ChR2-mediated effects in individual neurons, synaptic communication was inhibited by applying the ionotropic glutamate receptor antagonist kynurenic acid and the chloride channel blocker picrotoxin, which eliminates GABA_A_ receptor-mediated synaptic currents. Under such circumstances loose patch configuration - which spares the intracellular milieu of the recorded neurons - was conducted first to detect the light-evoked firing, followed by whole-cell recordings to measure the membrane responses upon light delivery (Fig. 2A). To reveal the firing characteristics of neurons, we also monitored the membrane potential responses upon intracellular injection of step currents with different amplitude (Fig. 2B). Biocytin content of intrapipette solution allowed us identifying unequivocally the recorded neurons *post hoc* (Fig. 2C). In these electrophysiological experiments we found that blue light illumination evoked spikes in most interneurons tested in addition to pyramidal neurons (Fig. 2A, C). Based on the firing features and neurochemical content of interneurons, light activation of ChR2 could discharge fast spiking PV-expressing interneurons, SST-containing interneurons showing accommodation in their firing and late spiking NPY-expressing neurogliaform cells in the hippocampus and mPFC (Supplementary fig. 1A-C). Moreover, there was a linear relationship between the number of spikes triggered by light delivery and the area of the light-evoked membrane voltage changes if the data for all recorded neurons were examined together (Fig. 2D). We also observed several neurons in which light illumination did not generate firing, yet there was a substantial membrane voltage response in them (see data points along the x axis), indicative for ChR2-mediated subthreshold responses. These data clearly demonstrate that blue light activation of ChR2 expressed under the control of CaMKIIα promoter can excite and effectively drive the firing of distinct types of interneurons, similarly to that observed in pyramidal neurons (Supplementary fig. 1D).

**Fig. 2.**
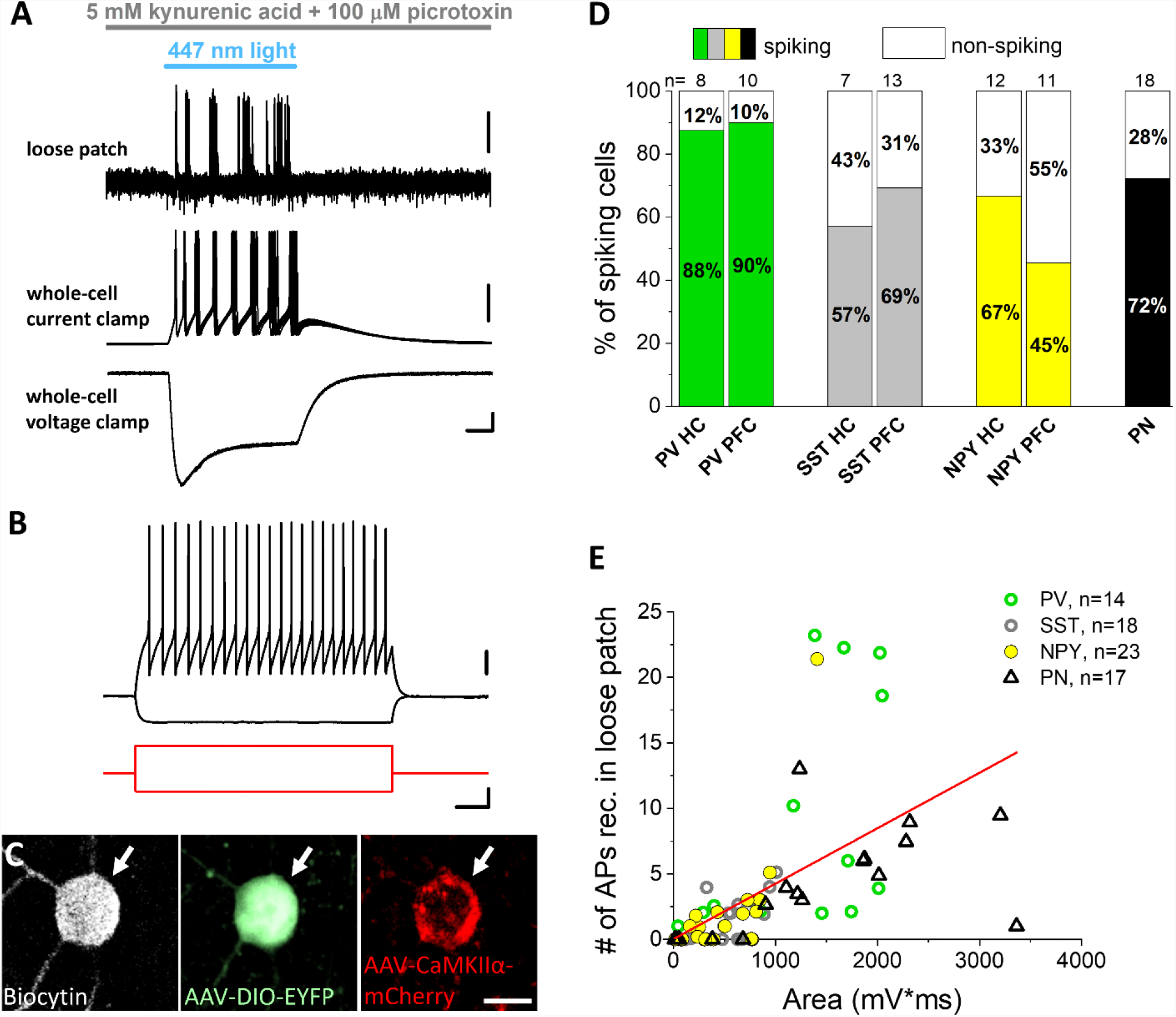
Blue light activation of Channelrhodopsin 2 (ChR2) expressed in parvalbumin (PV), somatostatin (SST) and neuropeptide Y (NPY)-containing interneurons under the control of CaMKIIα promoter readily evokes firing and direct voltage responses. **(A)** Ten consecutive, superimposed voltage traces from loose patch recordings (*top*) showing action potentials in response to ChR2 activation by blue light illumination in an inhibitory neuron sampled in an acute prefrontal cortical slice prepared from an Npy-Cre mouse. Representative voltage and current responses (*middle and bottom*) were recorded subsequently in the same interneuron in whole-cell mode as a result of ChR2 activation. Scale bars, loose patch, y=0.5 mV; current clamp, y=10 mV, voltage clamp, y=50 pA and x=10 ms. **(B)** Voltage responses of the interneuron with fast spiking phenotype (*top, black*) shown in (A) evoked by two current steps (*bottom, red*). Scale bars, top, y=20 mV; bottom, y=100 pA and x=100 ms. **(C)** Confocal images demonstrating that the recorded biocytin-filled inhibitory cell (white arrow) same as in (A) and (B) expresses Npy promoter-driven EYFP enhanced with anti-GFP immunostaining (*green, middle*) and CaMKIIα promoter-driven mCherry enhanced with anti-RFP immunostaining (*red, right*). Scale bar, 10 µm. **(D)** Ratio of spiking neurons upon light delivery monitored in loose patch experiments. **(E)** Area of ChR2-evoked voltage responses correlated significantly with the action potential number detected in loose patch mode during light illumination (Pearson’s r=0.56, p<0.001).

When the immunohistochemical data were compared with the results of slice physiology, there was a good correspondence, as all GABAergic cell types that expressed mCherry could also be discharged by light illumination, which agrees with their ChR2 expression. However, surprising exceptions were found as none but one PV-expressing and SST-expressing interneurons in the hippocampus and no SST-containing interneuron in the mPFC showed obvious signal for mCherry even after enhancement using immunostaining (Fig. 1B, Fig. 3), yet they readily spiked upon blue light delivery (Fig. 2C, Fig. 3). Therefore, we closely examined the mCherry content enhanced by immunostaining in fast spiking PV-containing interneurons and SST-containing interneurons that were depolarized upon light illumination. Surprisingly, we found only sparse immunolabeling for this reporter protein in spite of the large ChR2-medated voltage responses in fast spiking PV-containing interneurons sampled in the hippocampus and in SST-containing interneurons in the hippocampus and mPFC. For comparison, fast spiking PV-containing interneurons in the mPFC showed large light-evoked voltage responses as well as strong immunoreactivity for mCherry, in full agreement with our immunohistochemical results (Fig. 1B). In the BLA, mCherry content in SST-expressing interneurons varied (Fig. 3C). When the area of the voltage responses evoked by ChR2 activation was compared in interneurons with and without mCherry signal, a significant difference was observed (Fig. 3D). In spite of this difference in the postsynaptic response magnitude, the firing of interneurons having below-the-threshold mCherry labeling could be still evoked by blue light illumination (Fig. 2D, E; Fig. 3A-C). These unexpected observations indicate that light activation of ChR2 can affect the excitability even in those interneurons, where the presence of reporter proteins is ambiguous.

**Fig. 3.**
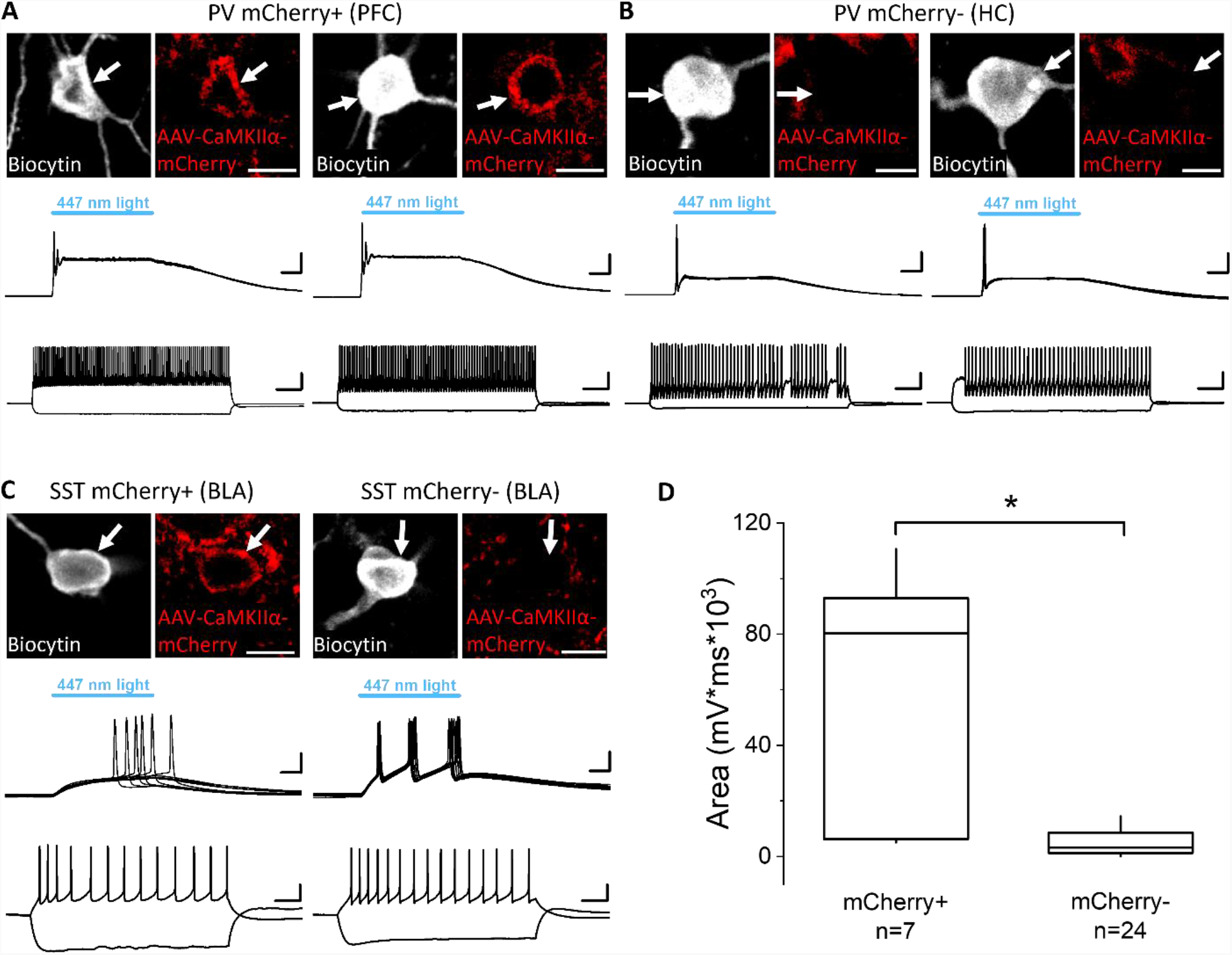
ChR2-evoked voltage responses that drive the firing in PV and SST interneurons can be detected in GABAergic cells with and without mCherry signal. **(A)** Two examples of mCherry-expressing PV interneurons recorded in the prefrontal cortex (PFC) filled with biocytin (*white, top*). mCherry signal was enhanced with anti-RFP immunostaining (*red, top*). Scale bars, 10 µm. Ten consecutive, superimposed voltage traces from whole-cell recordings (*middle*) showing voltage responses upon ChR2 activation by blue light illumination. Scale bars, y=10 mV and x=10 ms. Voltage responses of the interneurons (*bottom*) evoked by two current steps (+400 and −100 pA). Scale bars, y=20 mV and x=100 ms. **(B)** Two example PV interneurons recorded in the hippocampus (HC) filled with biocytin (*white, top*) that show no obvious CaMKIIα promoter-driven expression of mCherry even after enhancing the fluorescent protein signal with anti-RFP immunostaining (*red, top*). Scale bar, 10 µm. Ten consecutive voltage traces from whole-cell recordings (*middle*) showing voltage responses upon ChR2 activation by blue light illumination. Scale bars, y=10 mV and x= 10 ms. Voltage responses of the interneurons (*bottom*) evoked by two current steps (left, +400 and −100 pA; right, +300 and −100 pA). Scale bars, y=20 mV and x=100 ms. **(C)** Two examples of SST interneurons recorded in the basolateral amygdala (BLA) filled with biocytin (*white, top*); one shows clear mCherry signal following anti-RFP immunostaining (*red, top-left*), while the other displays no detectable expression of mCherry even after immunostaining using anti-RFP antibody (*red, top-right*). Scale bars, 10 µm. Ten consecutive traces from whole-cell recordings (*middle*) are superimposed showing voltage responses in both SST interneurons during blue light illumination that activates ChR2. Scale bars, y=10 mV and x=10 ms. Voltage responses of the interneurons (*bottom*) evoked by two current steps (left: +40 and −100 pA; right: +40 and −100 pA). Scale bars, y=20 mV and x=100 ms. **(D)** The area under the curve of ChR2-evoked voltage responses recorded in whole-cell mode is significantly larger in interneurons expressing mCherry than in those where the fluorescent protein signal was not present even after immunostaining. Data for PV and SST interneurons were pooled. p<0.01, Mann-Whitney U Test

Viral vector-mediated delivery of constructs typically produces higher levels of protein expression in the targeted neurons than in offspring crossed by genetically modified mice (Haery *et al*., 2019). Therefore, in the final set of experiments, we examined whether cortical interneurons express GFP signal in CaMKIIα-GFP mice. Based on the previous experience, much less GABAergic cells are expected to contain GFP in this transgenic mouse line in comparison with virus infected cortical regions. In line with the prediction, we indeed found a substantially lower number of GFP-expressing inhibitory cells visualized by immunostaining in the hippocampus, mPFC and the BLA of CaMKIIα-GFP mice. The highest ratio of GFP-positive neurons was observed in SST-containing interneurons in the BLA (34%), about 10-14% of nNOS-and NPY-expressing cells and virtually no PV-containing interneurons expressed GFP in any of the investigated cortical regions (data not shown). These results show that CaMKIIα promoter-driven constructs delivered by AAVs can significantly boost the expression of proteins in GABAergic cells in comparison to other transgenic approaches.

## Discussion

The main findings of the current study are as follows: i) CaMKIIα promoter can drive protein expression in at least four different GABAergic cell types in three different cortical structures; ii) the ratio of infected inhibitory cells varies between these brain areas revealed by immunohistochemical analysis; iii) light activation of ChR2 that expression is controlled by CaMKIIα promoter reliably drives spiking in cortical inhibitory cells and iv) ChR2-mediated excitation can be still significant even in those neurons where the reporter protein level is below the detection threshold of immunostaining.

Prior to our systematic study a publications has already indicated that PV-expressing interneurons in the marmoset cerebral cortex can be infected by AAVs carrying a CaMKIIα promoter driven construct (Watakabe *et al*., 2015). More recently it has been reported that in addition to PV interneurons, SST interneurons could also be targeted in the mouse motor cortex by CaMKIIα promoter using micoRNA guided tagging (Keaveney *et al*., 2020). Our findings are in agreement with these observations, but substantially advance those by showing that in addition to PV and SST inhibitory cells, GABAergic neurons expressing nNOS and NPY can be also a subject of infection by CaMKIIα promoter controlled constructs.

At present it is unclear why CaMKIIα promoter is able to drive protein expression in interneurons where neither immunolabeling, nor the sensitive RNAscope method revealed the presence of this protein kinase (Liu & Jones, 1996; Sik *et al*., 1998; He *et al*., 2021). Although CaMKIIα isoform is lacking from cortical GABAergic cells, another isoform, γCaMKII seems to be specifically expressed in inhibitory cells both in the hippocampus and neocortex (He *et al*., 2021). This observation raises the possibility whether the limited promoter of CaMKIIα using in constructs packed in AAV capsids may interact with the coding sequence of γCaMKII, which allows the protein expression in GABAergic cells using CaMKIIα promoter.

One of our interesting findings was the observation that hippocampal PV-containing interneurons as well as SST-expressing interneurons in the mPFC and hippocampus showed negligible or no immunoreactivity for mCherry, yet they expressed ChR2 at a level that was enough to reliably discharge them upon light delivery. These results point out an important concern, namely that the ambiguous visual detection of the reporter protein expression in neurons infected by AAV carrying constructs does not necessarily mean that the effector protein is not expressed at the level which can alter the function of reporter protein “lacking” neurons.

In summary, our study uncovered that in sharp contrast to the widespread belief, CaMKIIα promoter-driven expression of constructs is not specific for cortical glutamatergic neurons, but many types of GABAergic cells can be also infected. These results show the limitation of the use of CaMKIIα promoter in circuit studies aiming to target selectively the excitatory principal neurons in cortical structures and challenging the interpretation of previous studies. This constrain, however, can be overcome by the use of Vglut1-Cre mice, an approach that has been successfully applied to target selectively excitatory principal neurons, but not inhibitory cells first in the BLA (Andrasi *et al*., 2017), followed by subsequent studies in other cortical regions (Yamawaki *et al*., 2019; Luo *et al*., 2020).

## Materials and Methods

### Experimental animals

All experiments were approved by the Committee for the Scientific Ethics of Animal Research (PE/EA/131-4/2017, 993-7/2020) and were performed according to the guidelines of the institutional ethical code and the Hungarian Act of Animal Care and Experimentation (1998; XXVIII, section 243/1998, renewed in 40/2013) in accordance with the European Directive 86/609/CEE and modified according to the Directives 2010/63/EU. Every effort was taken to minimize the number of animals used and animal suffering. Three lines of transgenic mice (> 8 weeks old) expressing green fluorescent protein (GFP) under the control of the Pvalb promoter (BAC_Pvalb-GFP, (Meyer *et al*., 2002)), or expressing Cre recombinase under the Sst promoter (Sst-IRES-Cre, Jax.org, # 013044), or expressing Cre recombinase under the Npy promoter (BAC_Npy-Cre, MMRRC_034810-UCD), were used in *in vitro* experiments.

### Virus injections

For functionally testing ChR2 expression in GABAergic neurons Sst-Cre and Npy-Cre mice were injected with a mix of two different virus constructs, a channelrhodopsin (ChR2) and red fluorescent protein (mCherry) carrying virus (AAV2/5 CaMKIIα-ChR2(H134R)-mCherry-WPRE-hGH), and a Cre-dependent yellow fluorescent protein (EYFP) carrying virus (AAV1-EF1a-DIO - EYFP) in order to simultaneously label GABAergic interneurons. PV-GFP transgenic mice were injected only with the ChR2 and mCherry carrying virus (AAV2/5 CaMKIIα-ChR2(H134R)-mCherry-WPRE-hGH), because in this transgenic mouse line PV interneurons express GFP inherently. Viral vectors were obtained from Addgene. For injections the following volumes and stereotaxic coordinates were used (in mm): BLA (unilateral, right; 300 nl) AP: −1.8, ML: −3.0, DV: −3.7; dorsal hippocampus (unilateral, right; 2×200 nl) AP: −1.8, ML1: −1.5, ML2: −2.3, DV1: −1.3, DV2: −1.5; mPFC (bilateral; 200-200 nl) AP: +1.8, ML: 0.3, DV: 1.5.

### Immunohistochemical identification of virally labeled CaMKIIα-expressing neurons

P80-104 male C57Bl6/6J wild type mice were injected unilaterally with AAV 2/-CaMKIIα-ChR2(H134R)-mCherry-WPRE-hGH at three coordinates (in mm, to Bregma) aiming the mPFC (200 nl, AP +0.18, ML 0.3, DV −0.15), hippocampus (200 nl, AP −0.15, ML −0.15, DV −0.16) and BLA (200 nl, AP −0.15, ML −0.32, DV −0.44). After 6 weeks mice were anesthetized and transcardially perfused with 4% PFA, then the brain was removed and cut into 50 µm sections with a vibratome (Leica VT1000S). Sections ipsilateral to the injection side were incubated in a mixture of the following primary antibodies: rat-RFP (1:1,000, Chromotek), rabbit anti-somatostatin (1:10,000, Peninsula), guinea pig anti-parvalbumin (1:10,000, SYSY), goat anti-neuronal nitric oxide synthase (1:1,000, Abcam), revealed with the secondary antibodies (all 1:500, Jackson), Cy3-conjugated donkey anti-rat, Alexa 647-conjugated donkey-anti rabbit, Alexa488-conjugated donkey-anti guinea pig, Alexa405-conjugated donkey-anti goat. Another set of sections was incubated in a mixture of the following primary antibodies: rat-RFP (1:1,000, Chromotek), rabbit anti-somatostatin (1:10,000, Peninsula), guinea pig anti-neuropeptide Y (1:1,000, SYSY), goat anti-neuronal nitric oxide synthase (1:1,000, Abcam), revealed with the secondary antibodies (all 1:500, Jackson), Cy3-conjugated donkey anti-rat, Alexa 647-conjugated donkey-anti rabbit, Alexa 405-conjugated donkey-anti guinea pig and Alexa 488-conjugated donkey-anti goat. Sections were mounted in Vectashield (Vector laboratories) and 3D images were obtained using a Nikon C2 confocal microscope with Nikon CFI Super Plan Apo 20X objective (N.A. 0.75; z step size: 2 µm, xy: 0.31 µm/pixel). Colocalization of neurochemical markers in the neurons was examined and quantified manually with the Neurolucida 10.53 software (MBF Bioscience) and plotted with OriginPro 9.8 (OriginLab Corporation).

### Slice preparation and electrophysiological recordings

For *in vitro* experiments, acute brain slices containing the BLA and dorsal hippocampus or containing the mPFC were prepared. Mice >4 weeks post injection were decapitated under deep isoflurane anesthesia and the brain was quickly removed and placed into ice-cold cutting solution, containing (in mM): 252 sucrose, 2.5 KCl, 26 NaHCO_3_, 0.5 CaCl_2_, 5 MgCl_2_, 1.25 NaH_2_PO_4_, 10 glucose, bubbled with 95% O_2_/5% CO_2_(carbogen gas). Coronal slices of 200 μm thickness were prepared with a Leica VT1200S vibratome and transferred to an interface-type holding chamber filled with artificial cerebrospinal fluid (ACSF) containing (in mM): 126 NaCl, 2.5 KCl, 1.25 NaH_2_PO_4_, 2 MgCl_2_, 2 CaCl_2_, 26 NaHCO_3_, and 10 glucose, bubbled with carbogen gas. After incubation of the slices at 36°C for 60 minutes, slices were kept at room temperature until using them for recording. After at least 1 hour-long incubation, slices were transferred to a submerged type recording chamber perfused with ACSF bubbled with carbogen gas, additionally containing 5 mM kynurenic acid and 100 µM picrotoxin, at approximately 2-2.5 ml/min flow rate and 32°C.

Recordings were performed under visual guidance using differential interference contrast microscopy (DIC, Nikon FN-1) under 40x water dipping objective. Neurons expressing fluorescent protein either GFP or EYFP were visualized with epifluorescent illumination optics and detected with a CCD camera (Andor Zyla, NIS D software). Patch pipettes (4-7 MΩ) for whole-cell and loose patch recordings were pulled from borosilicate capillaries with inner filament (thin walled, OD 1.5 mm) using a P1000 pipette puller (Sutter Instruments). For loose-patch recordings pipettes were filled with ACSF, and an incomplete seal was formed during the recording with the cell membrane of the targeted neuron in order to monitor spiking activity in response to blue light (447 nm) stimulation. For whole-cell recordings the patch pipette contained the following (in mM): 110 K-gluconate, 4 NaCl, 2 Mg-ATP, 20 HEPES, 0.1 EGTA, 0.3 GTP (sodium salt), 10 phosphocreatine and 0.2% biocytin adjusted to pH 7.25 using KOH, with an osmolarity of 300 mOsm/L.

Recordings were performed with a Multiclamp 700B amplifier (Molecular Devices), low-pass filtered at 2 kHz, digitized at 10-25 kHz, and recorded with Clampex 10.4 (Molecular Devices). In loose patch recordings holding current was set to zero, in whole-cell mode, cells were held at the membrane potential of −65 mV. Recordings were analyzed with Clampfit 11.3 (Molecular Devices) and statistics were calculated and plotted in OriginPro 2021. Recordings were not corrected for junction potential.

To reveal the firing characteristics, neurons were injected with 800-ms-long alternating hyperpolarizing and depolarizing square current pulses with increasing amplitudes from −100 to 600 pA. Accommodating firing pattern, wide AP half-width and slow after-hyperpolarization were characteristic for control principal neurons. Principal neuron identity was further confirmed by the *post hoc* morphological analysis of their spiny dendrites and morphology.

To test the ChR2 expression, interneurons containing GFP or EYFP were recorded in loose patch mode while laser light pulses (447 nm) were applied. The length of the laser light pulse was set to 50-ms and at maximum intensity in order to elicit firing in neurons with slow membrane kinetics as well (Andrasi *et al*., 2017). To prevent activation of the recorded cell by network effect and to prevent contamination of ChR2 responses by synaptic events, the superfused ACSF solution contained ionotropic glutamate and chloride channel blockers (5 mM kynurenic acid and 100 µM picrotoxin). Fluorescent neurons that were firing action potentials during light stimulation were considered as responding cells, while in non-responding cells no action potential could be elicited with light stimulation. Ratio of responding/non-responding neurons were calculated from the total number of tested fluorescent cells in loose patch experiments. In order to define the ChR2 expression level in the recorded cells, whole-cell patch clamp recordings were made in the same cell immediately after loose patch recording using a different pipette containing K^+^-based intrapipette solution. Light evoked ChR2 current and voltage responses were recorded in voltage clamp and current clamp mode, respectively. To define the functional ChR2 level in GABAergic cells that is high enough to make these neurons responsive to light stimulation, voltage responses were plotted against the number of action potentials in response to light stimulation. Area under the curve was calculated from average voltage response of a cell in the time window during light stimulation. As a control of CaMKIIα expression, random neighboring principal cells were recorded both in loose patch and whole-cell mode identically to interneuron recordings. Pearson’s correlation coefficient was calculated for revealing relationship between the excitability of neurons in loose patch mode and voltage changes monitored in whole-cell mode in response to ChR2 activation. To assess whether mCherry+ and mCherry-PV and SST interneurons at the injection site have a difference in the level of ChR2 expression, area under the curve of light-evoked voltage responses obtained by whole-cell voltage clamp recordings were compared. The significant difference was assessed with Mann-Whitney U Test.

### Immunohistochemistry on slices following electrophysiological recordings

After *in vitro* recordings slices were fixed in 4% paraformaldehyde overnight at 4°C and biocytin content of the recorded neurons was visualized using Alexa 647-conjugated streptavidin (1:10,000 or 1:20,000, Molecular Probes). mCherry and GFP or EYFP signals in the neurons were enhanced by immunostainings using the following antibodies: rat anti-RFP (1:1,000, Chromotek) revealed by Cy3-conjugated donkey anti-rat secondary antibody (1:500, Jackson) and chicken anti-GFP (1:1,000, SYSY) revealed by Alex 488-conjugated donkey anti-chicken antibody (1:500, Jackson). Slices were mounted in Vectashield (Vector laboratories) and 3D images were obtained using a Nikon C2 confocal microscope with Nikon CFI Super Plan Apo 20X objective (N.A. 0.75; z step size: 1 µm, xy: 0.31 µm/pixel). Images were processed and analyzed with the NIS Elements AR 5.30.01 software (Nikon Instruments).

### Immunohistochemical identification of CaMKIIα-expressing neurons in a transgenic mouse line

Two male CaMKIIα/GFP-FVB/AntF1 mice (P53-67)(Wang *et al*., 2013) was anesthetized and transcardially perfused with 4% PFA, then the brains were removed and cut into 50 µm sections with a vibratome (Leica VT1000S). Sections were incubated in a mixture of the following primary antibodies: chicken anti-GFP (1:1,000, SYSY), rabbit anti-somatostatin (1:10,000, Peninsula), guinea pig anti-parvalbumin (1:10,000, SYSY), goat anti-neuronal nitric oxide synthase (1:1,000, Abcam). Primary antibodies were revealed either with the secondary antibodies (all 1:500, Jackson): Alexa 488-conjugated donkey-anti chicken, Alexa 647-conjugated donkey-anti rabbit, Cy3-conjugated donkey anti-guinea pig; or with Alexa 488-conjugated donkey-anti chicken, Alexa 647-conjugated donkey-anti rabbit and Cy3-conjugated donkey anti-goat. Another set of sections was immunostained with chicken anti-GFP (1:1,000, SYSY) revealed with Alexa 488-conjugated donkey-anti chicken and guinea pig anti-neuropeptide Y (1:1,000, SYSY) revealed with Cy3-conjugated donkey-anti guinea pig secondary antibody. Sections were mounted in Vectashield (Vector laboratories) and 3D images were obtained using a Nikon C2 confocal microscope with Nikon CFI Super Plan Apo 20X objective (N.A. 0.75; z step size: 2 µm, xy: 0.31 µm/pixel). Colocalization of the neurochemical markers in the neurons was examined and quantified manually with the Neurolucida 10.53 software (MBF Bioscience).

## Acknowledgements

We acknowledge financial support from the Hungarian Brain Research Program (2017-1.2.1-NKP-2017-00002) awarded to NH. The authors are grateful to Éva Krizsán and Erzsébet Gregori for their excellent technical assistance. We also thank László Barna, the Nikon Microscopy Center at the Institute of Experimental Medicine, Nikon Austria GmbH, and Auro-Science Consulting, Ltd., for kindly providing microscopy support.

The authors declare no competing financial interest.

## Author contributions

Designed experiments: JMV, TA, NH

Performed experiments: JMV, TA, PNP

Analyzed data: JMV, TA, PNP

Supervised the project: JMV, NH

Wrote the paper: JMV, TA, NH

---

**Supplementary figure 1 to Fig. 2.**
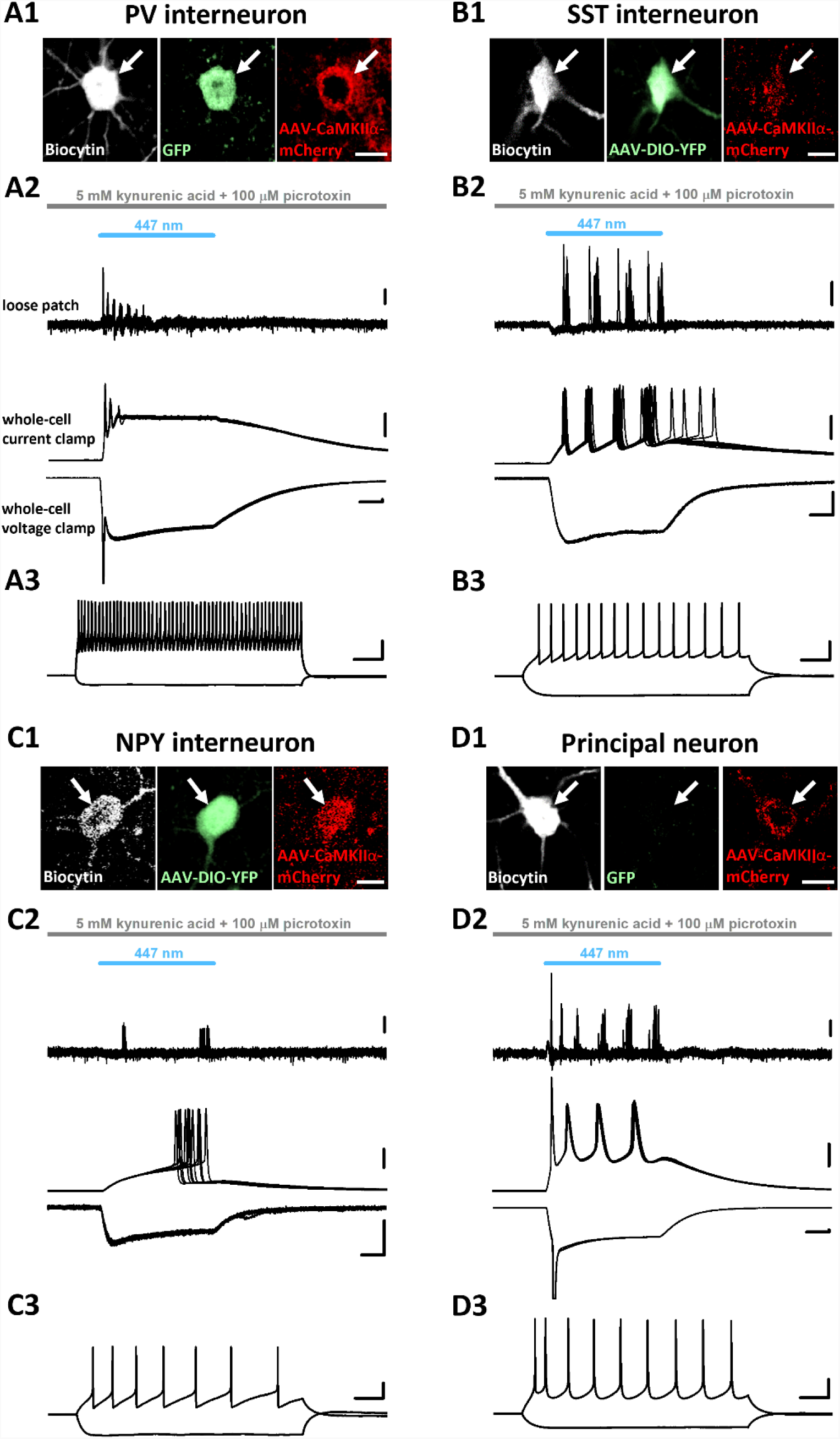
Example recordings from different cortical regions show that all three examined GABAergic cell types and principal neurons were successfully infected with AAVs carrying CaMKIIα-ChR2-mCherry construct. (A1-D1) Confocal images demonstrating that the recorded inhibitory cells and a principal neuron filled with biocytin (*white, left*) express mCherry under the control of CaMKIIα promoter. The signal of the red fluorescent protein was enhanced with anti-RFP immunostaining (*red, right*). Biocytin-filled GABAergic cells expressed GFP or EYFP under the Pvalb, Sst and Npy promoters, respectively, and its fluorescent protein content was enhanced *post hoc* with anti-GFP immunostaining (*green, middle*) (A1-C1). In the principal neuron no GFP/EYFP signal (*green, middle*) was detected (D1). Scale bar, 10 µm. **(A2-D2)** Ten consecutive voltage traces are superimposed from loose patch recordings (*top*) showing action potentials in response to ChR2 activation by blue light illumination in a PV, SST, NPY interneuron and a principal neuron (same as shown in A1-D1), respectively. Example neurons were sampled in the prefrontal cortex, except the SST interneuron, which was recorded in the hippocampus. Representative voltage and current responses (*middle and bottom*) were recorded subsequently in the same PV, SST, NPY interneurons and principal neuron in whole-cell mode as a result of ChR2 activation. Scale bars, loose patch, 0.2 mV; current clamp, 20 mV; voltage clamp, y=100 pA and x=10 ms. **(A3-D3)** Voltage responses of the example neurons (*top, black*) evoked by two current steps (A3: +500 and −100 pA; B3: +200 and −100 pA; C3: +50 and −90 pA; D3: +150 and −100 pA). Scales, y=20 mV and x=100 ms.

